# Hybrid Deep Learning and Lee-Carter Model for Mortality Forecasting: A Study of US Adults Aged 35-80

**DOI:** 10.1101/2025.10.13.682203

**Authors:** Collins Abaitey, Eric Abakah, Clement Asare, Emmanuel Oppong, Fareeda Maltiti Baba-Sadiq, Elshaddai Appiah

## Abstract

This study introduces a hybrid approach that enhances mortality forecasts by integrating machine learning techniques specifically Long Short-Term Memory (LSTM), Bidirectional LSTM (BiLSTM), and Neural Networks (NN) with the traditional Lee-Carter model. After deriving the time index *κ*_*t*_ from the Lee-Carter model, these deep learning models were employed to capture complex temporal patterns, which were then incorporated into the Lee-Carter framework to improve forecasting accuracy. The hybrid models were evaluated using historical mortality data from the United States, covering ages 35 to 80 years from 1975 to 2020. Among the models tested, the LSTM model outperformed all others, demonstrating superior capability in capturing time dependencies and producing more accurate mortality forecasts. The integration of LSTM with the Lee-Carter framework led to significant improvements in predictive accuracy. This research demonstrates that combining traditional statistical approaches with modern deep learning techniques, particularly LSTM, offers a powerful method for enhancing mortality forecasting by effectively modeling time-dependent patterns. These findings provide valuable insights and tools for policymakers, actuaries, and healthcare professionals to improve planning and decision-making.

## 1. INTRODUCTION

Hybrid mortality modeling has become increasingly relevant as researchers strive to improve the accuracy and reliability of mortality forecasts. In actuarial science, demography, and public health, accurate mortality predictions are crucial for understanding and preparing for future trends in life expectancy and mortality rates. Hybrid approaches, which integrate traditional statistical methods with machine learning techniques, offer enhanced capabilities for capturing complex patterns in mortality data. This integration allows for more precise forecasting, accommodating both short-term fluctuations and long-term trends in mortality rates.

The importance of accurate mortality forecasting is highlighted by its significant economic implications. For example, life insurance companies and pension funds face considerable vulnerability to longevity risk, which arises when policyholders live longer than anticipated, resulting in the need for higher-than-expected disbursements. The global market for longevity risk is substantial, with pension-related liabilities alone estimated to be between USD 60 trillion and USD 80 trillion [2]. According to the World Health Organization, global life expectancy increased significantly from 66.8 years in 2000 to 73.4 years in 2019, representing a gain of more than six years [31]. Although healthy life expectancy increased from 58 years to 64 years during this period, reflecting an 8% rise, this improvement has been mainly attributed to reductions in mortality rates rather than a decrease in years lived with disability. In other words, the overall increase in life expectancy (6.6 years) has outpaced the growth in healthy life expectancy (5.4 years). This decline in mortality rates across all age groups can largely be attributed to significant advancements in nutrition, public health, and medical research aimed at the prevention and treatment of diseases. Furthermore, the COVID-19 pandemic highlighted the urgency of accurate mortality prediction. During the pandemic, the United States alone saw life insurance companies pay over ninety billion dollars to beneficiaries due to COVID-19-related deaths [1]. This demonstrates the critical role of mortality forecasts in managing unexpected health crises and economic risks.

There is a long history of mortality modeling that began with the publication of Gompertz’s law of mortality in his seminal paper [9]. Since then, numerous models have been proposed, although the endeavor to forecast mortality remains relatively recent. Currently, many of the mortality forecasting techniques in use, particularly those employed by European statistical offices, are extrapolative in nature [28]. These methods leverage observed consistency in age patterns and mortality trends over time, as noted in [4] and cited in [14]. Extrapolative techniques are considered more objective and user-friendly, and they are often thought to yield more accurate forecasts compared to expectation methods—based on expert opinions—and explanation methods, which predict mortality by specific causes or explanatory models [4].

In 1992, Lee and Carter [18] introduced a novel approach to long-term mortality forecasting, known as the Lee-Carter (LC) model. For many years, the Lee and Carter [18] methodology has been the standard extrapolative technique for mortality projections. This model decomposes mortality into two components: a time-varying effect that causes all age-specific central mortality rates to exhibit a consistent pattern of stochastic evolution over time, and two age-specific parameters for each age group. However, despite its remarkable capabilities and robust performance, the Lee-Carter model has several limitations as highlighted by several researchers [17, 20, 22, 11, 25]. Notably, its assumption of constant age-specific mortality reduction rates leads to unrealistic estimates and underestimation of mortality improvements among older populations [6].

In recent years, hybrid forecasting models that incorporate deep learning and other machine learning approaches have gained increasing significance across various fields, including actuarial science and computer science. Several studies have highlighted the importance of these models [20, 22, 11]. The key emphasis is about how machine learning techniques offer a more flexible and data-driven method for predicting mortality trends. That is, Machine learning abilities to capture intricate interconnections and nonlinear relationships between covariates and mortality outcomes. Additionally, machine learning approaches can also enhance the evaluation of mortality estimates produced by traditional stochastic models, such as the Lee-Carter and Renshaw-Haberman models [7]. There is a growing body of innovative research focused on employing machine learning methods in both the non-life and life sectors, particularly in the realm of mortality forecasting [26, 7]. However, some previous studies have combined the Lee-Carter model with fixed effect regression models such as random forest and decision trees [17]. Nonetheless, these models often fail to adequately represent time-dependent patterns and nonlinear correlations in mortality data, resulting in erroneous estimates [11]. In actuarial and demographic applications, neglecting to include temporal dependence in mortality trends can lead to poor risk management and decision-making. Therefore, to enhance the accuracy of mortality forecasts and capture the complex time-dependent patterns in mortality data, this study investigates the use of deep learning models to predict *κ*_*t*_ in the Lee-Carter model.

The primary aim of this study is to hybridize the Lee-Carter model with deep learning algorithms, such as Long Short-Term Memory (LSTM), Bidirectional LSTM (Bi-LSTM), and neural networks, to enhance the accuracy of predicting time-dependent mortality trends. The objectives of the study are 1) to develop a hybrid mortality forecasting model by integrating the Lee-Carter model with deep learning techniques such as LSTM, Bi-LSTM, and neural networks, and 2) to evaluate the performance of these deep learning models in predicting mortality for individuals aged 35 to 80. Achieving these objectives will answer the following research questions: 1) How can the integration of the Lee-Carter model with deep learning techniques improve the accuracy of mortality predictions for the target age group? 2) How do the predictive performances of the deep learning models LSTM, Bi-LSTM, and neural networks compare in terms of accuracy and reliability? This study is significant as it seeks to improve mortality forecasting accuracy, particularly for the aging population and the associated financial risks of longevity. By integrating traditional statistical methods with advanced deep learning techniques, this research aims to enhance predictive capabilities essential for policymakers, public health officials, and insurance providers. The Focus on individuals aged 35 to 80 years also provides insights into this demographic’s unique health challenges. Ultimately, the findings could inform effective mortality management strategies and policy decisions, contributing to a better understanding of contemporary mortality dynamics. The rest of the study is structured as follows: Section 2 presents the materials and methods; Section 3 presents the results with discussions; and Section 4 concludes the study with recommendations for future research.

## 2. MATERIALS AND METHODS

### Dataset

This study analyzed secondary data from the Human Mortality Database for a selected sub-population of the United States aged 35 to 80 from 1975 to 2020. The dataset includes a total of 89,139,773 individuals, comprising 49,202,432 females and 39,937,341 males. Within this population, there were 1,257,755 recorded deaths, with 509,970 females and 747,785 males. The data encompasses several variables: year, age, group (gender), mortality, exposure, and deaths. To analyze mortality trends over time in the United States, we computed the annual log mortality rate using the following formula:

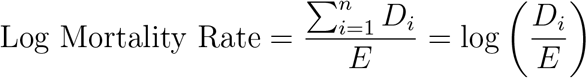

where *D*_*i*_ represents the total number of deaths for age group *i* in a specific year and *E* is the total population at risk (exposure) in that age group. Examining these age-specific mortality rates over several years allowed for a better understanding of fluctuations in mortality risk both within and between age groups.

Figure 1 illustrates the relationship between age and mortality rate. The plot reveals a clear upward trend, indicating that individual mortality rates increase with age. This consistent rise in mortality rates underscores the critical role of age in understanding and forecasting mortality trends. Such a pattern is anticipated and aligns with the fundamental premise of many mortality forecasting models, which assert that age is the primary determinant of mortality probability [30, 29]. Additionally, there is a pronounced increase in mortality rates among individuals aged 70 and older. This steep rise highlights the heightened vulnerability of older age groups to mortality, emphasizing the need for targeted interventions and healthcare strategies tailored to this population.

**Figure 1.**
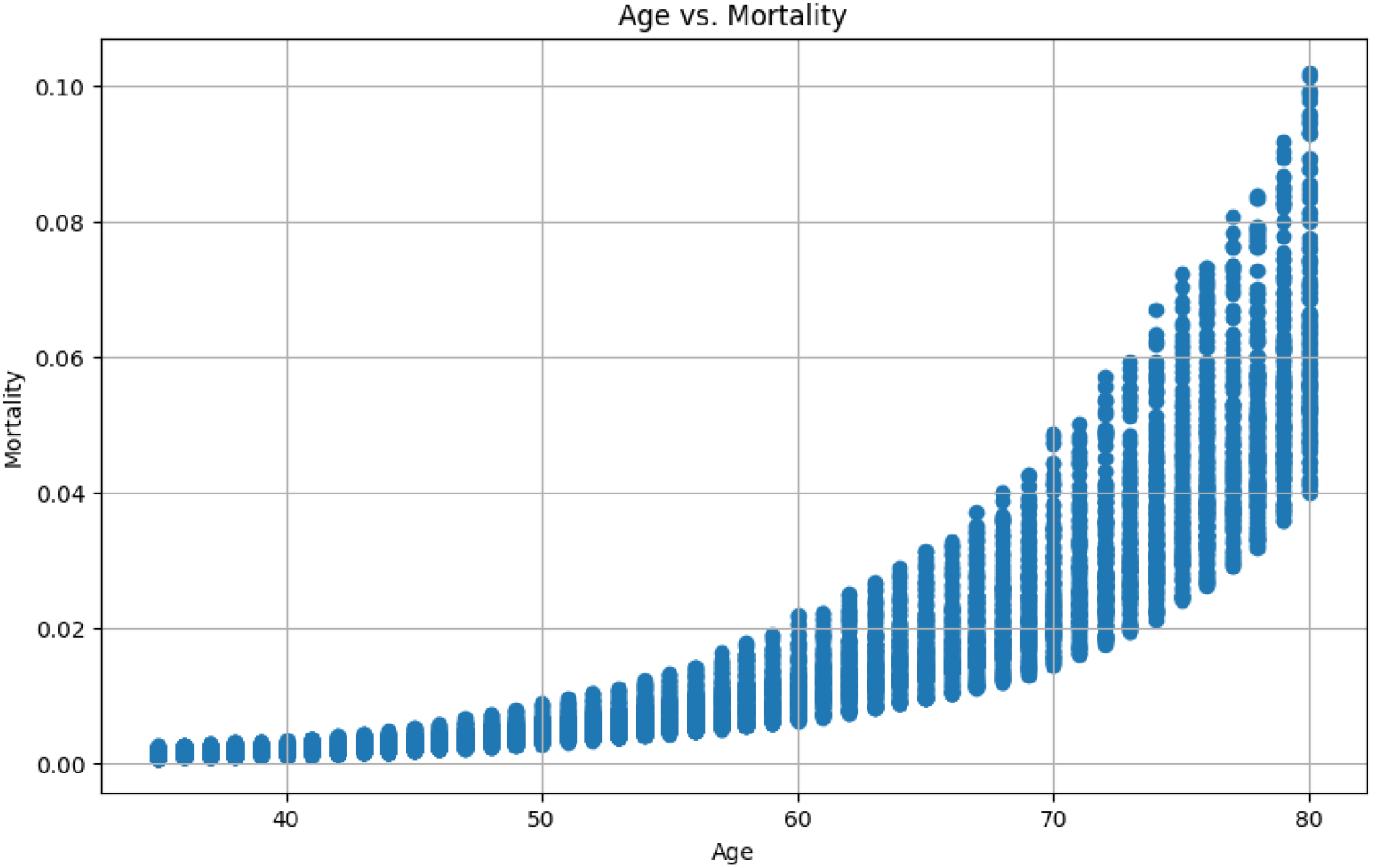
Plot of age against mortality rate

**Figure 2.**
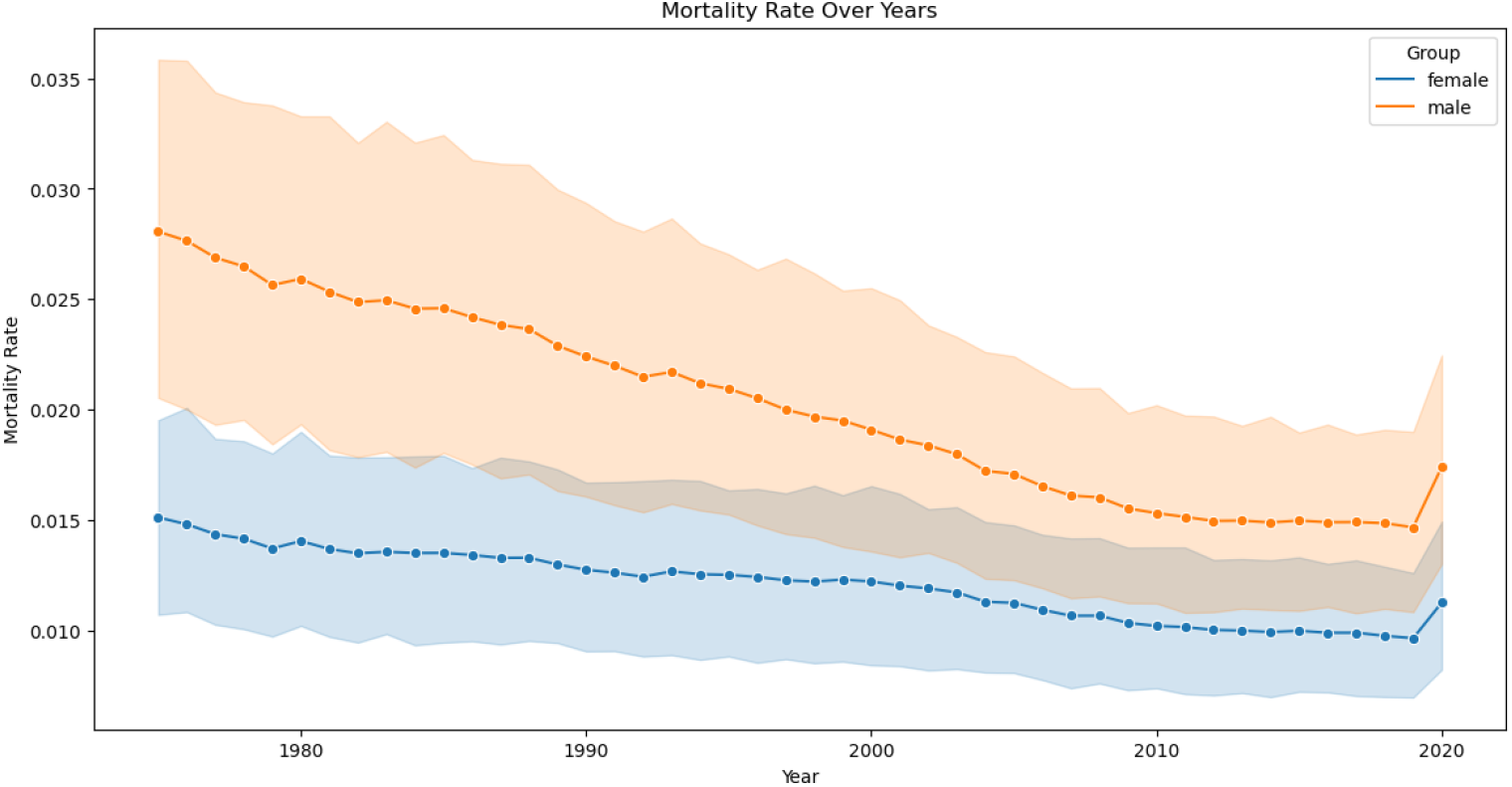
Plot of mortality rates pattern against years.

Figure 3 illustrates a general decline in mortality rates for both females and males over the years, reflecting an overall improvement in mortality trends. A notable feature of the graph is the persistent gap between the blue and orange lines, which indicates that males consistently experience higher mortality rates than females. This observation aligns with established patterns across various populations, where males exhibit a 60% greater mortality risk compared to females [32]. However, the gradual reduction of this disparity over time suggests a trend toward convergence, indicating that gender differences in mortality may be diminishing. Interestingly, the most recent data points, clustered around 2020, reveal a marked increase in mortality rates for both sexes. This upward trend can be linked to the significant impact of the COVID-19 pandemic, which has profoundly affected mortality patterns across the globe.

**Figure 3.**
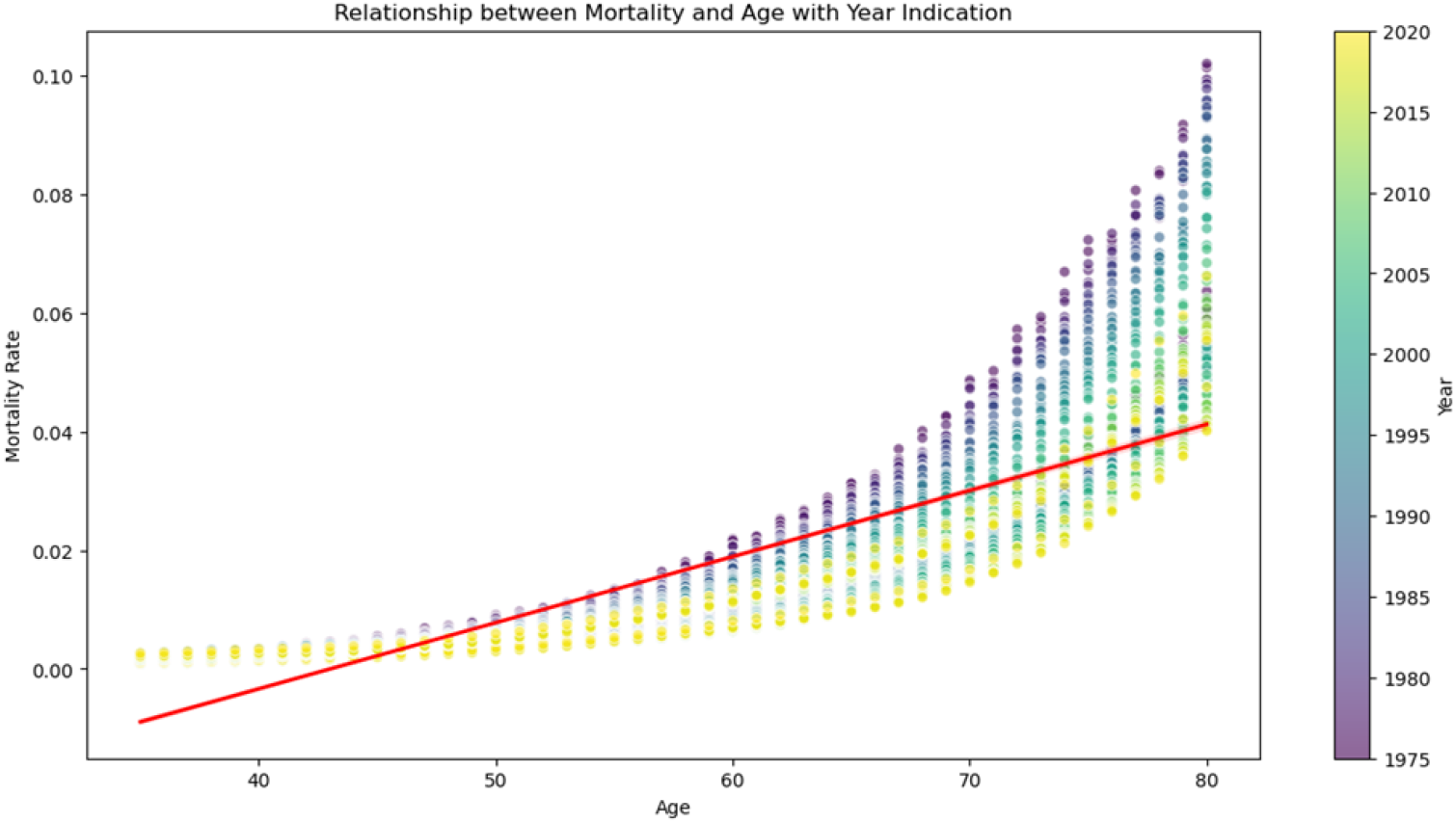
Plot of mortality rates pattern by age group.

### Methods

We develop a hybrid model that integrates the traditional Lee–Carter model with the three deep learning algorithms (LSTM, BiLSTM, and NN). We seek to hybridize the Lee-Carter model with each of the three deep learning models. This allows for the creation of distinct hybrid models, enabling us to evaluate the performance of each hybrid approach in forecasting mortality rates.

### Lee-Carter Model

One popular stochastic model for predicting mortality rates is the Lee-Carter model. In the fields of actuarial sciences and demography, it has grown in importance as a technique for predicting and projecting age-specific death rates. The foundation of the Lee-Carter model is the notion that the overall level of mortality is represented by a time-varying index (t = 1,., T) and the log of the mortality rate at a specific age (x = 1,., X) which captures the distinct age pattern of death. The Lee-Carter model can be represented mathematically as follows.

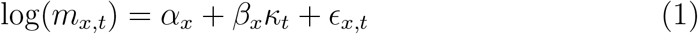

where *m*_*x,t*_ is the central rate of mortality for age group x at time t. The *α*_*x*_ parameter is the age-specific component of the average age pattern of mortality which shows how mortality rates vary across different ages. The *β*_*x*_ parameter measures the sensitivity of mortality rates to changes over time for each age group. It shows how changes in the overall mortality level affect mortality rates at different ages concerning the time-dependence. The *κ*_*t*_ parameter is a time-varying parameter that characterizes the overall mortality level in a given year or time. It is interpreted as a measure of the average mortality risk at time t. The *ϵ*_*x,t*_ (0, *σ*^2^) is the error term that accounts for the unobserved factors and random variability in mortality rates for each age group *x* and time t.

To ensure that equation 1 can be identified, then: Σ*β* = 1, Σ *κ* = 0. To estimate equation 1, we employ a Singular Value Decomposition (SVD) approach described in [18]. We then estimate the age-specific intercept term by finding the average logarithmic transformed death rate for each age over time.

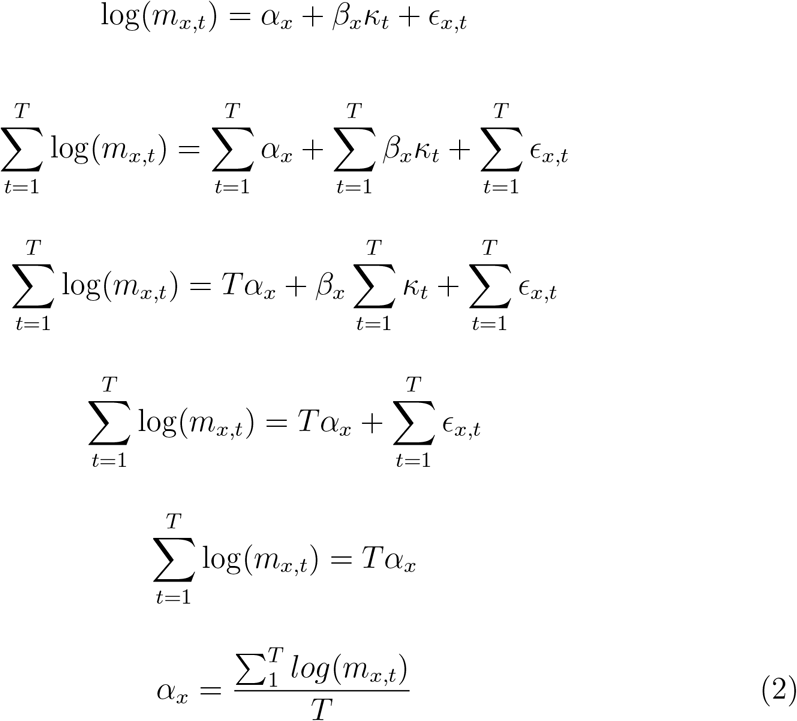

We Subtract these averages from the log-transformed rates to center the data:

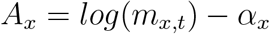

The matrix A consists of these centered values, where A is an *XT* matrix (with X being the number of age groups and T being the number of periods). We apply the SVD to the matrix of logarithms of rates after subtracting the log of the age-specific rates. We continue by modeling the time index *κ*_*t*_ to minimize deviations in the logs of the mortality rates.

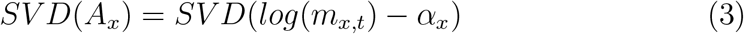

The result of Equation 3 will be three matrices: U, ∑, and *V* ^*T*^ where U is an orthogonal matrix where the value depends on age, *V* ^*T*^ is an orthogonal matrix where its value is time-dependent and is a singular diagonal matrix.

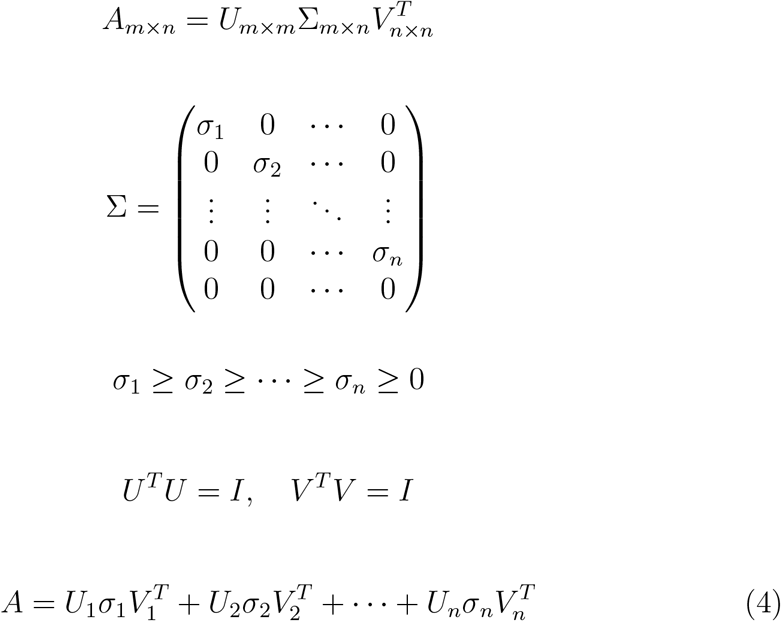

We used a rank of 1 for the estimation of the parameters *β*_*x*_ and *κ*_*t*_. *β*_*x*_ is the first column of U and *κ*_*t*_ is the first column of V.

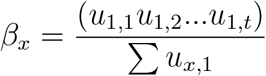

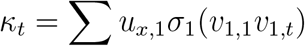

### Long Short-Term Memory Model

LSTM is a form of RNN that was created to address the limitations of standard RNNs in capturing long-term dependencies in sequential data. It was introduced in [12]. The key principle behind how LSTM works is that, rather than using the same feedback loop connection for events that occurred long ago and events that occurred yesterday to forecast what will happen tomorrow, LSTM employs a more sophisticated structure incorporating gates to regulate information flow. Specifically, LSTM networks use a forget gate, an input gate, and an output gate to regulate which information is retained, updated, and sent on to the next time step. This design enables LSTM to efficiently preserve and use both long-term and short-term memories. This allows LSTMs to perform language-related tasks.

At each time step *t* of the LSTM Layer Computations, the Long Short-Term Memory (LSTM) layer performs the following operations:

The forget gate determines which information from the previous cell state should be retained and carried forward to the next cell state. The activation of the forget gate *f*_*n*_ is computed as:

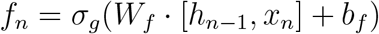

where *σ*_*g*_ is the sigmoid activation function, *W*_*f*_ is the weight matrix for the forget gate, [*h*_*n*−1_, *x*_*n*_] represents the concatenation of the previous hidden state *h*_*n*−1_ and the current input *x*_*n*_, and *b*_*f*_ is the bias term. The input gate decides how much of the new information should be added to the current cell state. The activation of the input gate *i*_*n*_ is computed as:

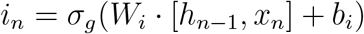

Here, *W*_*i*_ is the weight matrix for the input gate, and *b*_*i*_ is the bias term. The temporary cell state 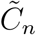 represents the new candidate values for the cell state, which are scaled by the input gate before being added to the current cell state. It is computed as:

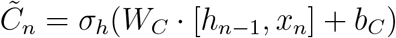

where *σ*_*h*_ is typically the hyperbolic tangent (tanh) activation function, *W*_*C*_ is the weight matrix for the cell state, and *b*_*C*_ is the bias term. The current cell state *C*_*n*_ is updated by combining the previous cell state, scaled by the forget gate, and the new candidate values, scaled by the input gate:

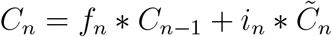

The forget gate determines how much of the previous cell state *C*_*n*−1_ to keep, while the input gate determines how much of the new candidate state 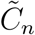 to add. The output gate determines the output of the LSTM cell. The activation of the output gate *o*_*n*_ is computed as:

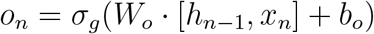

where *W*_*o*_ is the weight matrix for the output gate, and *b*_*o*_ is the bias term. The hidden state *h*_*n*_ is the output of the LSTM cell at the current time step. It is computed by applying the output gate to the tanh of the current cell state *C*_*n*_:

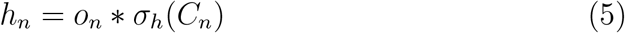

LSTM uses two key activation functions, the sigmoid function transforms any input into a value between 0 and 1. It is defined as:

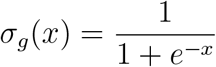

The tanh function transforms any input into a value between -1 and 1. It is defined as:

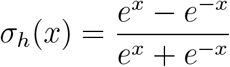

### Bidirectional Long Short-Term Memory Model

Bidirectional LSTMs are a variant of the standard two-long short-term memory (LSTM) that is also designed to handle sequential data. The main idea behind Bi-LSTMs is to analyze the input sequence both forwards and backward in order to allow the model to incorporate information from both past and future states.

The Bi-LSTM architecture is made up of two independent LSTM layers, one of which processes the input sequence forward and the other backward. The outputs of these two LSTM layers are then merged at each time step to get the final result. Unlike standard RNNs, which process input sequences in just one way (either forward or backward), Bi-LSTM processes the sequence in both directions concurrently. The forward LSTM layer is the same as the LSTM model defined in Equation 5. Mathematically, the backward LSTM layer computes the following equations at each time step *t*:

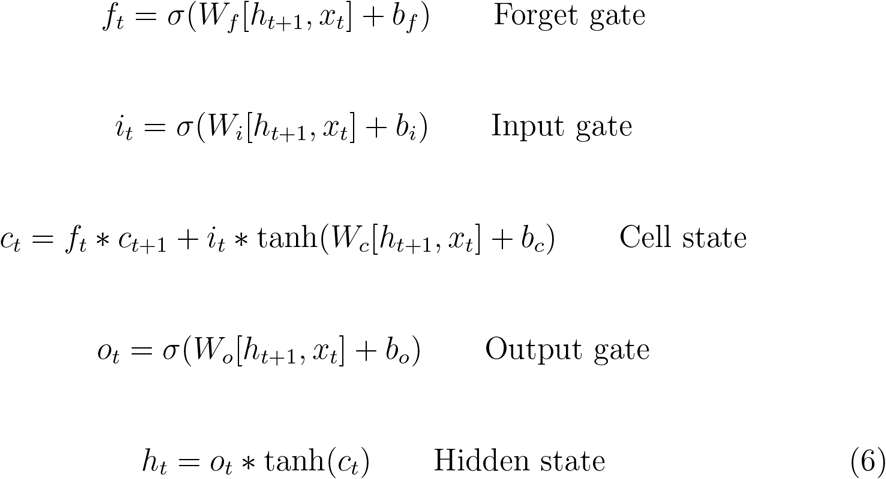

During the forward pass, the input sequence moves through the LSTM layer from the beginning to the end. At each step, the forward LSTM calculates its state and adjusts its memory cell using the current input along, with the previous hidden state and memory cell. Meanwhile, the input sequence is also processed by the LSTM layer in order starting from the last step and going back to the first. Like in the forward pass the backward LSTM computes its state and updates its memory cell based on the present input and previous hidden state and memory cell.

After both forward and backward passes are finished the hidden states from both LSTM layers are merged at every step. This merging can be as simple as concatenating the hidden states or applying some other transformation. One advantage of Bi-LSTM is its ability to capture not the preceding context, at a given time step (unlike RNNs) but also the subsequent context. By taking into account both future details Bi-LSTM can grasp intricate relationships, within the input sequence.

### Neural Network Model

Neural networks are powerful machine learning models inspired by biological neural networks, capable of learning complex patterns and representations from data, [10]. It is composed of three layer: input layer, hidden layers and output layer which are connected by neurons. The neurons within a neural network receive inputs and compute a weighted sum, followed by the application of an activation function that introduces non-linearity. During the training phase, the network undergoes iterative adjustments of connection weights and biases. This process aims to minimize the disparity between predicted and actual outcomes, a technique known as optimization. The neural network is trained using a supervised learning approach, where the model learns to map input data to the desired output by minimizing a loss function, [5].

The activation of a neuron *j* in layer *l* is computed as:

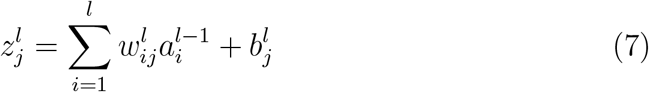

where 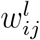 is the weight, 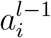 is the activation from the previous layer, and 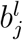 is the bias term. In this study, we An activation function *σ* is then applied to compute the final activation:

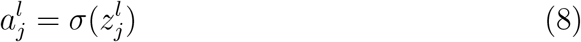

The output layer produces the final prediction, and the performance of the model is measured using a loss function such as the Mean Squared Error (MSE):

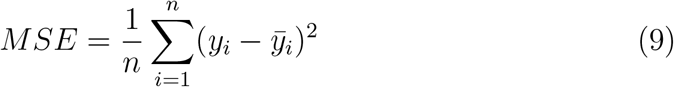

where *y*_*i*_ is the actual value and 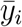 is the predicted value.

Neural networks are trained using optimization methods such as Gradient Descent and backpropagation, which modify the weights to reduce the loss function. This allows neural networks to capture complex trends, making them crucial for tasks like credible mortality forecasting.

### Formation of the Hybrid Mortality Forecasting Model

We lagged the data before using it to train the LSTM, Bi-LSTM, and neural network models. Lagging the data captures the temporal dependencies inherent in time series data which allows the models to learn from past values and generate accurate future predictions. Lagged data creates a sequence for the model to learn from, allowing it to identify trends, seasonality, and temporal patterns [19, 3]. We assumed that the first three years’ *κ*_*t*_ values could serve as a reliable indicator for forecasting the fourth year’s value. This approach from [22] has shown to be an excellent method of estimating the time series variable *κ*_*t*_, which is also described by [13].

The following table illustrates the dataset used for the LSTM, Bi-LSTM, and NN models, including the actual *κ*_*t*_ values and their lagged versions:

**Table 1:** *κ*_*t*_ values used for training and testing the models.

| Table 1: $\kappa_t$ values used for training and testing the models | | | | |
| --- | --- | --- | --- | --- |
| Year | $\kappa_t$ values | lag1 | lag2 | lag3 |
| 1978-01-01 | 9.605133 | 10.171352 | 11.318423 | 12.040781 |
| 1979-01-01 | 8.279512 | 9.605133 | 10.171352 | 11.318423 |
| 1980-01-01 | 8.768251 | 8.279512 | 9.605133 | 10.171352 |
| 1981-01-01 | 7.834674 | 8.768251 | 8.279512 | 9.605133 |
| 1982-01-01 | 7.133978 | 7.834674 | 8.768251 | 8.279512 |
| ⋮ | ⋮ | ⋮ | ⋮ | ⋮ |
| 2016-01-01 | -9.022713 | -8.882640 | -9.100307 | -9.005688 |
| 2017-01-01 | -9.040511 | -9.022713 | -8.882640 | -9.100307 |
| 2018-01-01 | -9.326591 | -9.040511 | -9.022713 | -8.882640 |
| 2019-01-01 | -9.794005 | -9.326591 | -9.040511 | -9.022713 |
| 2020-01-01 | -3.408362 | -9.794005 | -9.326591 | -9.040511 |

In the study, we employed an 80%-20% data split to allow for a thorough evaluation of our models’ capabilities, with 80% of the data used for training and the remaining 20% serving as the testing dataset. We used the lagged data for training, while the actual *κ*_*t*_ values were used for testing.

For the initial forecasts, we used LSTM, Bi-LSTM, and Neural Networks. Each model was trained on the lagged training dataset, and 5-fold crossvalidation was used to assess and evaluate the models’ prediction performance while avoiding overfitting. Using cross-validation, we were able to successfully integrate the predictions from each model. The hybrid model properly predicts the time index *κ*_*t*_ for the Lee-Carter model. This integration improves the Lee-Carter model’s ability to forecast mortality rates. The forecast values are then substituted into the original Lee-Carter model to estimate mortality rates. The age-specific parameters in the Lee-Carter model are expected to remain constant across time.

To assess the predictive accuracy of our proposed model on data, we generated predictions for each individual model, particularly LSTM, Bi-LSTM, and NN. We next compared three widely utilized Root Mean Squared Error (RMSE), Mean Absolute Error (MAE), and Mean Absolute Percentage Error (MAPE) to assess the performance of each model. Our goal was to obtain lower values of RMSE, MAE, and MAPE for our suggested models, since this suggests improved prediction performance and generalization to the data. The error metrics utilized in this study are mathematically defined as follows:

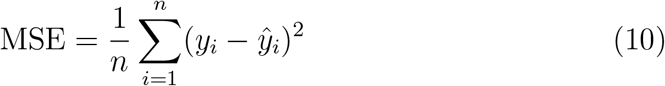

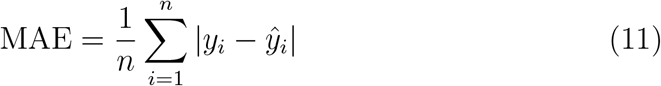

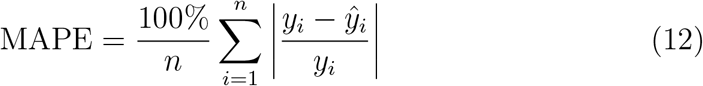

where *y*_*i*_ represents the actual values, ŷ_*i*_ denotes the predicted values, and *n* indicates the number of observations.

## 3. RESULTS AND DISCUSSION

### Results

Figure 3 shows the relationship between age and mortality rates, with data points color-coded by year and ranging from 1975 to 2020. The red regression line overlays the scatter plot, which emphasizes the overall trend between age and the mortality rate. The color gradient changes from purple (previous years) to yellow (recent years) to add a time component to the analysis. The figure clearly shows a positive correlation between age and mortality rates. As individuals age, their mortality rate steadily rises. The mortality rate increases gradually between the ages of 35 and 50. The scatter points are densely packed and relatively low, which implies a consistent and reduced mortality risk among younger people. This slow move implies that while mortality does increase with age, the danger is quite minor until the age of 50. It then starts to rise significantly between the ages of 50 and 65. The points begin to spread out further, and the slope of the regression line becomes steeper than in the 35-50 age range. After the age of 65, it accelerates rapidly. The points go up the y-axis, and the regression line’s steep slope indicates a considerable increase in mortality. The significant increase suggests that people over 65 have a significantly higher chance of death.

Figure 4 shows the mortality trends of individuals in the United States from age 35 to age 80 from the year 1975 to the year 2020. From the plot, we can see an overall downward trend in mortality rates for most age groups from 1980 to about the year 2010. This decreasing trend indicates that advancements in various areas, including healthcare, living conditions, and overall quality of life, have led to a progressive drop in mortality rates over the last few decades. However, this positive trend appears to be disrupted around the year 2020, with a notable increase in mortality rates across practically all age groups. This sudden rise could be attributed to the COVID-19 pandemic, which had a considerable influence on worldwide death rates, particularly among older age groups who are more vulnerable to the virus, [21].

**Figure 4.**
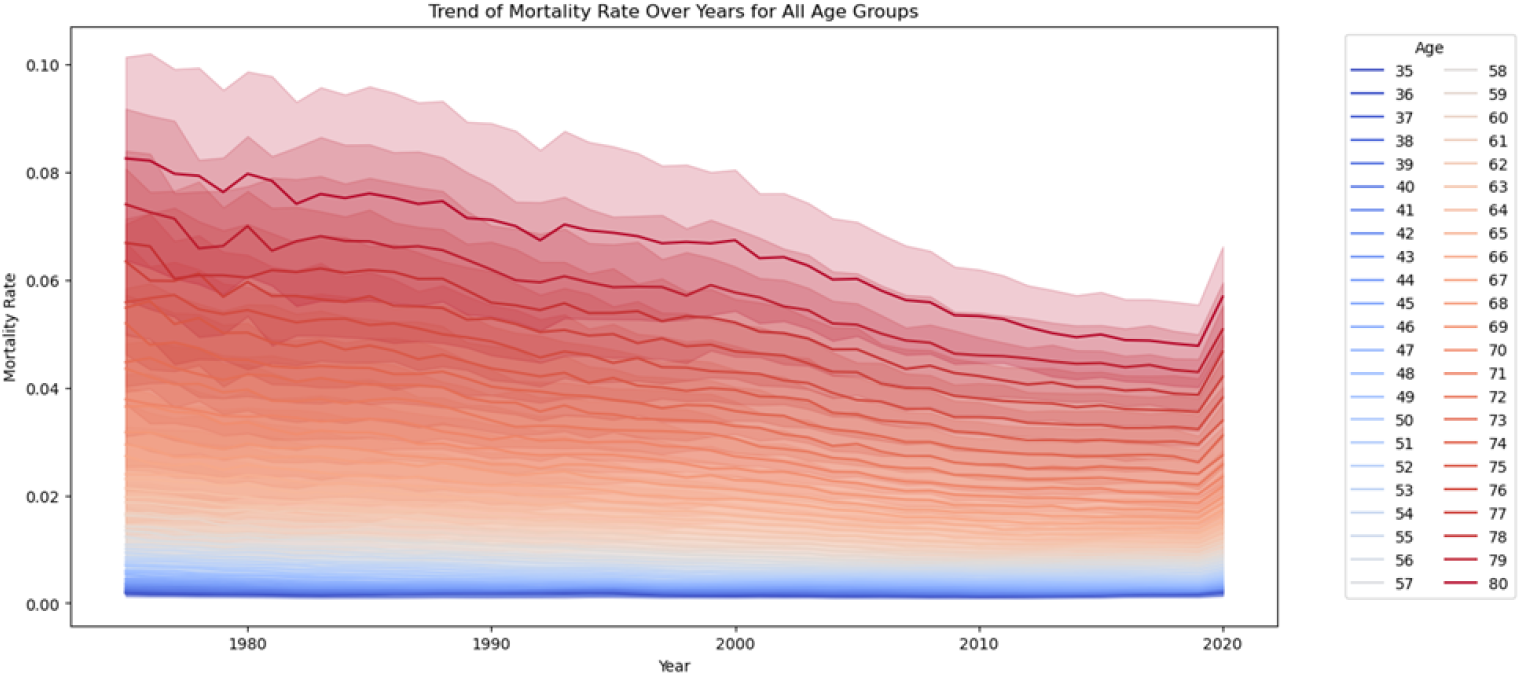
Plot of a trend showing mortality rate against years for all age groups

Tables 2, 3, 4 present the estimated *k*_*t*_ values for males, females, and the total population as derived from the Lee-Carter model using Singular Value Decomposition (SVD). The *k*_*t*_ values serve as indicators of mortality rates, which exhibit a general declining trend over time.

**Table 2:** Estimated male *κ*_*t*_.

| Year | kt_values |
| --- | --- |
| 1975 | 14.401631 |
| 1976 | 13.733420 |
| 1977 | 12.610189 |
| 1978 | 12.027303 |
| 1979 | 10.693669 |
| ⋮ | ⋮ |
| 2016 | -10.676321 |
| 2017 | -10.693316 |
| 2018 | -10.777154 |
| 2019 | -11.297666 |
| 2020 | -4.767075 |

**Table 3:**
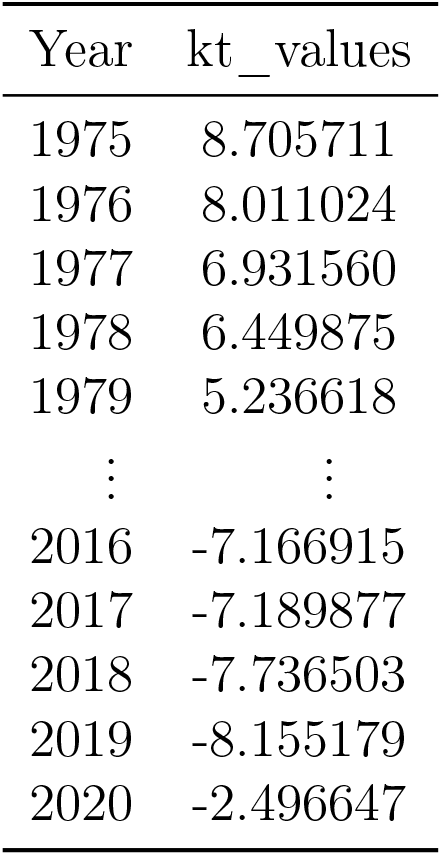
Estimated female *κ*_*t*_.

| Year | kt_values |
| --- | --- |
| 1975 | 8.705711 |
| 1976 | 8.011024 |
| 1977 | 6.931560 |
| 1978 | 6.449875 |
| 1979 | 5.236618 |
| ⋮ | ⋮ |
| 2016 | -7.166915 |
| 2017 | -7.189877 |
| 2018 | -7.736503 |
| 2019 | -8.155179 |
| 2020 | -2.496647 |

**Table 4:** Estimated Total *κ*_*t*_ (1975 - 2020)

| 1975-1997 |  | 1998-2020 |  |
| --- | --- | --- | --- |
| Year | $\kappa_t$ | Year | $\kappa_t$ |
| 1975 | 12.04078149 | 1998 | -0.05833998 |
| 1976 | 11.31842258 | 1999 | -0.20085416 |
| 1977 | 10.17135173 | 2000 | -0.80713630 |
| 1978 | 9.60513269 | 2001 | -1.51180769 |
| 1979 | 8.27951194 | 2002 | -1.98819315 |
| 1980 | 8.76825079 | 2003 | -2.68310507 |
| 1981 | 7.83467391 | 2004 | -4.23448208 |
| 1982 | 7.13397824 | 2005 | -4.47961205 |
| 1983 | 7.18130481 | 2006 | -5.63225902 |
| 1984 | 6.76811536 | 2007 | -6.60085383 |
| 1985 | 6.74531104 | 2008 | -6.71262441 |
| 1986 | 6.25808518 | 2009 | -7.79976763 |
| 1987 | 5.75464300 | 2010 | -8.43229879 |
| 1988 | 5.56548467 | 2011 | -8.70511172 |
| 1989 | 4.51438524 | 2012 | -9.07272625 |
| 1990 | 3.72790227 | 2013 | -9.00568829 |
| 1991 | 3.18708345 | 2014 | -9.10030728 |
| 1992 | 2.49613971 | 2015 | -8.88263957 |
| 1993 | 2.99694232 | 2016 | -9.02271288 |
| 1994 | 2.33911521 | 2017 | -9.04051145 |
| 1995 | 2.03341385 | 2018 | -9.32659108 |
| 1996 | 1.33569908 | 2019 | -9.79400455 |
| 1997 | 0.44426087 | 2020 | -3.40836221 |

Figure 5 presents the estimated *β*_*x*_ values derived from the Singular Value Decomposition (SVD), plotted against age groups measured in years. The graph illustrates that different age groups exhibit varying degrees of change in mortality rates, as reflected in their corresponding *β*_*x*_ values. Notably, the rate of change is most pronounced for individuals aged 35 to 38, with a slight decline observed after age 38. Subsequently, there is a significant increase around age 40, followed by another decline at age 63. This comprehensive depiction enhances our understanding of the dynamic patterns in mortality rate variations across various age groups over the observed period.

**Figure 5.**
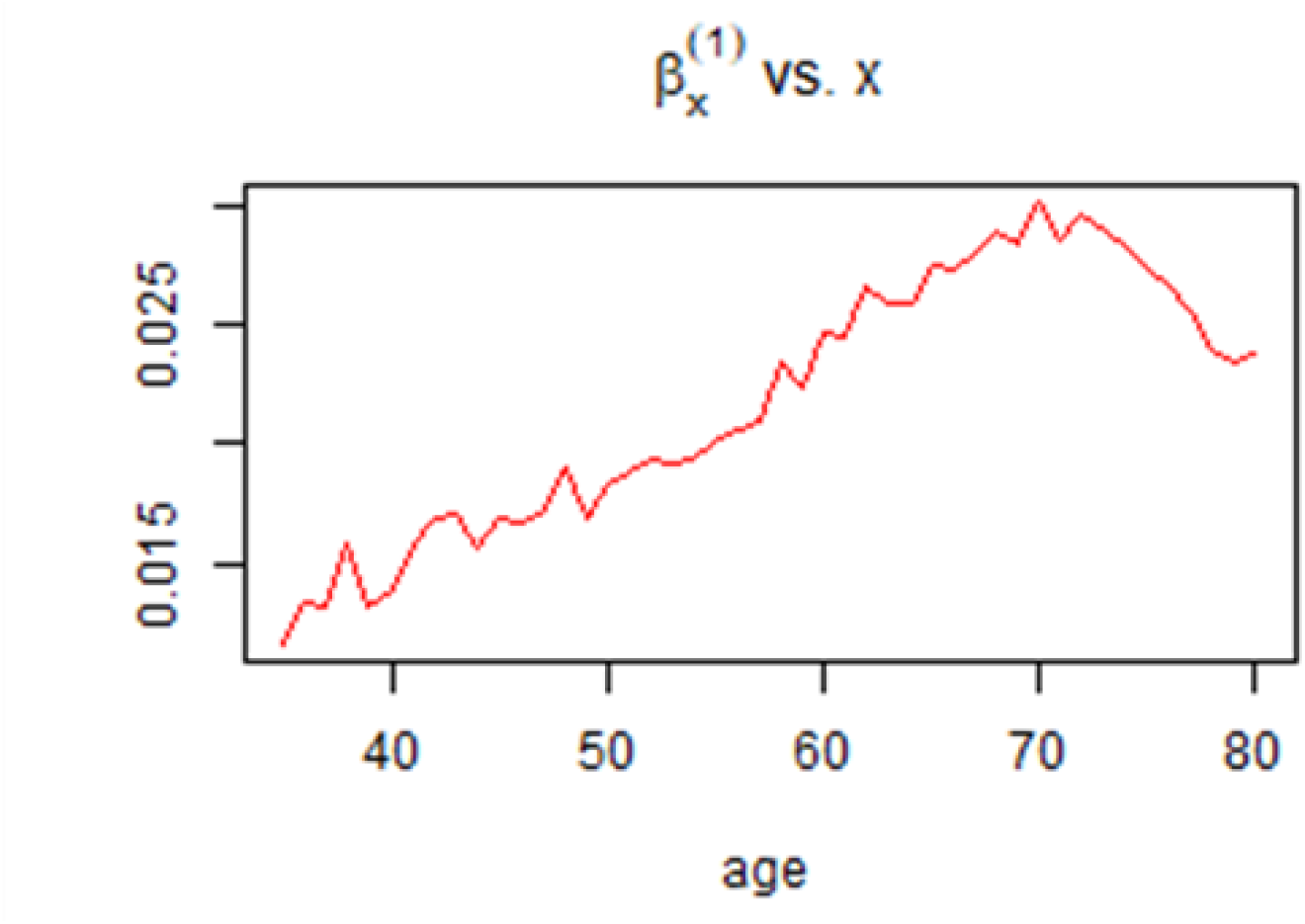
Plot of *β*_*x*_ of the Lee-Carter model against age

Analyzing these plots allows for a clearer understanding of how mortality rates are structured across different ages and how they evolve over time for females. The trends depicted in these parameters facilitate more accurate forecasting of future mortality rates by capturing the underlying patterns and their progression.

### Application of the LSTM, Bi-LSTM, and the neural network models to the United States mortality data set

In this section, we delve into the performance evaluation of three machine learning models: Long Short-Term Memory, Bidirectional LSTM, and Neural Network. The evaluation procedure focuses on assessing each model’s predicted accuracy, using evaluation metrics that give essential insights into their respective performances. We summarize the error metrics in the Table 5.

**Table 5:** Error Metrics.

| Metric | MSE | MAE | MAPE |
| --- | --- | --- | --- |
| LSTM | 5.3590 | **1.2734** | **0.3083** |
| Bi-LSTM | **5.3298** | 1.2973 | 0.3101 |
| NN | 5.4586 | 1.4879 | 0.3292 |

From the Table 5, we can observe that the Bi-LSTM has the lowest MSE indicating that it has the smallest average squared errors. The LSTM tends to have the lowest MAE and MAPE indicating that on average, LSTM’s predictions are closer to the actual values. With the lowest MAE and MAPE, the LSTM showed superior in predicting *κ*_*t*_.

We compare the predicted *κ*_*t*_ values from 2015 to 2019, produced using hybrid models, with the actual *κ*_*t*_ values. The predicted *κ*_*t*_ values are then used to estimate mortality rates by replacing them into the original Lee Carter equation. We assume constant values for *α*_*x*_ and *β*_*x*_, as proposed by Lee and Carter in 1992. Table 6 shows predicted and actual death rates for the years 2015 to 2019.

**Table 6:** Actual and Predicted *κ*_*t*_ Values.

| Year | Actual $\kappa_t$ | Predicted $\kappa_t$<br>(LSTM) | Predicted $\kappa_t$<br>(Bi-LSTM) | Predicted $\kappa_t$<br>(NN) |
| --- | --- | --- | --- | --- |
| 2015 | -8.8826396 | -8.803983 | -8.768156 | -8.543645 |
| 2016 | -9.0227129 | -8.776583 | -8.737502 | -8.488410 |
| 2017 | -9.0405115 | -8.773879 | -8.737778 | -8.438983 |
| 2018 | -9.3265911 | -8.780485 | -8.749879 | -8.477440 |
| 2019 | -9.7940046 | -8.842212 | -8.811502 | -8.616756 |

### Discussion

Our mortality forecasting study identified persistent patterns in mortality rates, with older individuals experiencing a higher mortality rate compared to younger individuals. This finding aligns with previous research by Pierce et al. [23], which revealed that mortality rates rise exponentially with age. Contributing factors include weakened immune systems, accumulated health risks over time, and increased susceptibility to chronic diseases [8]. These results underscore the necessity for specialized healthcare treatments tailored to distinct age groups to address their unique vulnerabilities. While preventive measures and early detection programs may be more effective for older populations, it is crucial for younger individuals to address lifestyle and environmental factors to mitigate long-term health risks.

Our analysis of mortality rates in the US revealed a consistent decline across all age groups over time. This positive trend can be attributed to several factors, including improved education, better healthcare services, and enhanced living conditions [33]. Increased health awareness and access to medical facilities have led to earlier disease identification and treatment, significantly reducing mortality rates. Public health initiatives, such as immunization programs and anti-infective disease campaigns, have been instrumental in this decline. Furthermore, improved nutrition and hygiene have yielded broad health benefits by reducing the incidence of malnutrition and waterborne infections. This integrated approach to health has resulted in a more robust population capable of effectively managing diseases.

The time index *k*_*t*_ from the Lee-Carter model demonstrates a strong decreasing trend, indicating a systematic decline in mortality rates. This pattern suggests improvements in population health and lifespan, consistent with the demographic transition hypothesis [24]. This hypothesis explains how a country’s birth and death rates shift from high to low as it develops. Factors such as better healthcare, improved living conditions, and public health initiatives contribute to this decline [15]. Advances in medical technology, enhanced access to healthcare services, and effective disease prevention and management programs have all played a role in this upward trend. These enhancements not only increase life expectancy but also improve the quality of life for many individuals.

Initially, the error metrics for our models were significant, reflecting challenges in effectively collecting and modeling historical data. Machine learning models, including LSTM, Bi-LSTM, and neural networks, are particularly sensitive to data quality issues such as noise and outliers [34]. These challenges can lead to overfitting or underfitting, adversely affecting the forecast accuracy of each model. Our findings are consistent with other research on mortality prediction. For instance, Scognamiglio et al. [27] demonstrated that several recurrent neural network models, including LSTM and Bi-LSTM, frequently outperform traditional models like the Lee-Carter model in capturing nonlinear patterns and providing accurate projections. These neural network models excel at processing time-series data and can learn complex temporal relationships, making them well-suited for mortality forecasting. However, the study also highlighted the challenges of predicting sudden fluctuations in mortality rates, a common issue across all modeling methodologies [16]. In our case, the unexpected occurrence of the COVID-19 pandemic caused a sudden shift in mortality rates, complicating accurate forecasts.

## 4. CONCLUSION

This study aimed to enhance mortality forecasting by integrating the LeeCarter model with hybrid forecasting models, including LSTM, Bi-LSTM, and Neural Networks, focusing on accurately capturing temporal dependencies in mortality trends. Utilizing lagged data for training enabled effective modeling of these trends, with the LSTM hybrid model outperforming the Bi-LSTM and Neural Networks due to its capacity to capture long-term dependencies in sequential data. While Bi-LSTM models may lead to overfitting, the LSTM’s emphasis on historical data points proved more effective for mortality forecasting. The LSTM-Lee-Carter hybrid model yielded the most accurate mortality rate estimates, illustrating the potential of combining hybrid models with traditional actuarial frameworks to improve prediction accuracy. The findings offer valuable insights for policymakers, actuaries, and healthcare practitioners, enhancing decision-making and preparation for demographic and health-related challenges.

